# Transcriptomic study of testicular hypoxia adaptation in Tibetan sheep

**DOI:** 10.1101/2024.06.20.599912

**Authors:** Binyan Yu, Yanan Yang, Yijian Li, Rong Gao, Min Ma, Xinrong Wang

## Abstract

The Tibetan sheep is a typical hypoxia-tolerant mammal, which lives on the plateau, an altitude of between 2,500 and 5,000 meters above sea level, the study of its hypoxic adaptation mechanism provides reference for exploring the hypoxic adaptation mechanism of other animals. To grope for the genetic mechanism of adaptation to hypoxic environment at the transcriptional level in Tibetan sheep testicular tissue, and to identify candidate genes and key pathways related to sheep adaptation. Histological observation of testicular tissues from two sheep breeds was carried out using hematoxylin-eosin (HE) conventional staining. A total of 277 differentially expressed genes (DEGs) were authenticated at high altitude Tibetan sheep (ZYH) and low altitude Tibetan sheep (ZYM) by RNA sequencing technology (RNA-Seq), which included 165 up-regulated genes and 112 down-regulated genes. Functional analyses revealed several terms and pathways, such as were closely related with testis adaptation to plateau. Several genes (including *GGT5, AGTR2, EDN1, LPAR3, CYP2C19, IGFBP3, APOC3* and *PKC1*) were remarkably enriched in several pathways and terms, which may impact the Plateau adaptability of sheep by adjusting its reproductive activity and sexual maturation, and protecting Sertoli cells, various spermatocytes and spermatogenesis processes. The results make a reasonable case for a better understanding of the molecular mechanisms of adaptation to altitude in sheep.

## 1. Introduction

In the high altitude environment, although the living conditions are harsh, different species of animals have evolved specific adaptations (hypoxia resistance, cold, enhanced metabolic capacity) to better accommodate the high altitude environment, such as Tibetan pig[1], Himalayan marmot[2]and Tibetan migratory locusts[3]. In male mammals, the testis is a highly specialised tissue[4], whose function is to produce sperm and androgens[5]. Both processes are influenced by unfavorable plateau environmental conditions, like hypoxia, high ultraviolet rays and low temps. Hypoxia impaired the quality of sperm germ cells, caused an imbalance of sex hormones (testosterone and estradiol) and slowed testicular development[6]. Previous researches have revealed the effect of plateau hypoxia and hypothermia on semen quality, reproductive hormones, and spermatogenesis in humans[7], monkeys[8], mice[9]and rats[10].Therefore, it is crucial to study the regulatory genes that influence hypoxic adaptation in the testicular tissue.

As one of the most prevalent and widely distributed animals on the Plateau, Tibetan sheep has excellent adaptability to the plateau environment of low oxygen[11]. While this discrepancy in breed altitude adaptation may be due to differences in foraging or reproductive methods, differences due to genetic factors may also be a major aspect. accordingly, It is important for the sheep industry to understand the mechanisms behind the differences in testicular adaptation of Tibetan sheep at high and low altitudes. Studies have used RNA-seq to identify the transcriptomes of ovaries, lungs, and liver[12–14], which identified the high altitude adaptation-related genes.

Accordingly, in the study, comprehensive analysis of testicular transcriptome data in high and low altitude Tibetan sheep using RNA-Seq, to discover the molecular mechanism of plateau adaptation in Tibetan sheep and searching for genes and regulatory pathways related to plateau adaptation, in order to offer a theoretical justification for the production and reproduction of Tibetan sheep.

## 2. Material and Method

### 2.1 Ethical Statement

The study complied with the requirements of the Committee for the Keeping and Use of Live Research Animals in Gansu Province, China. The Ethics Committee of Gansu Agricultural University granted all animal laboratory procedures (approval number: 2019-044). The animals were never exposed to unnecessary pain.

### 2.2 Collection of animals and samples

Twelve healthy Tibetan sheep aged one year were chosen for this study, including six Tibetan sheep from Hualong, Qinghai province (average altitude 2200m) and six Tibetan sheep from Haiyan, Qinghai province (average altitude 3700m). All experimental individuals were given the same nutrition under the same feeding conditions and had free access to food and water. After the experimental animals were slaughtered, the collected testis samples were separated two parts, one part was promptly chilled with liquid nitrogen and lay in -80°C for future experiments, and the other tissue samples were fixed with 4% paraformaldehyde for histological analysis.

### 2.3 Preparation of testicular tissue sections and analysis of data

Fresh tissue blocks from Tibetan sheep at different elevations were rinsed with tap water, dehydrated, xylene clear, embedded in paraffin and cut into 5μm tissue sections. Testicular tissue sections were then dewaxed and stained with HE (hematoxylin and eosin), and fixed and sealed with neutral resin. With the Olympus DP 71 microscope (Olympus Optical Co., Ltd., Tokyo, Japan) observes and photographs the sections. Four slices were randomly selected, Five non-repeating horizons were randomly chosen for each slice at 200x and 400x resolutions, and relevant values were taken with software (Image Pro plus 6.0). Measurement data were represented as mean ± SD (standard deviation). Using the student’s t-test by SPSS software (version 27.0) to do statistically analyzed , and a P value of less than 0.05 was considered as statistically significant.

### 2.4 RNA Extraction, cDNA Library Preparation, and Illumina Sequencing

According to the manufacturer’s procedure, the specimens of Tibetan sheep testis at high and low elevation were collected. Trizol (TransGen Biotech, Beijing, China) was used to extract total RNA from the testis samples. The completeness of RNA samples was assessed using Agilent 2100 (Agilent Technologies, Santa Clara, CA, USA) and 260 / 280 OD values of 1.8∼2.0 samples were Chosen for further analysis to use, the eligible RNA was utilized for constructing libraries. Each sample required 1 µg of total RNA for library preparation. The Qubit 2.0 and Agilent 2100 were used to determine concentration and insert size of prepared libraries. The generated cDNA libraries were sequenced on the Illumina sequencing platform.

### 2.5 Quantification of gene expression and identification of DEGs

For the purpose of getting the clean reads, the raw data underwent filtration through the elimination of adapters, low mass sequences and poly-n-containing reads. Filtered reads alignments were Compared with reference genome by software (HISAT2)[15]. Matched reads were assembled using String Tie software and their gene expression was calculated[16]. Expression levels of gene were estimated as number of transcript fragments per kilobase / million fragment mapping reads (FPKM)[17]. The raw data collected in the study has been archived in NCBI’s SRA. Serial number for this project is PRJNA10718. Expression abundance of different treatment groups was determined using DEseq2 software to identify differentially expressed genes (DEGs), The differentially expressed genes were selected as: fold changes (FC)≥ 1.5, significance *P*<0.05.

### 2.6 GO and KEGG Functional Enrichment Analysis

To learn more about the function of DEGs in testicular of Tibetan sheep at high and low elevations, The study utilized the GO terms and the KEGG pathway to analyze the correlation between the DEGs and the various biological functions of genes, and the criterion of significant enrichment was a *P*<0.05

### 2.7 RT-qPCR for Validation of DEGs

Testing the accuracy of transcriptome sequencing data, ten genes were chosen for Quantitative real-time-PCR (*RT-*q*PCR*). By Oligo 7.0 and Primer 5.0 constructed Primers (Table 1). RNA samples were reverse transcribed into cDNA using a reverse transcriptase kit (Takara, Dalian, China). The *RT-*q*PCR* cycle parameters were as follows: three biological replicates were used in each experiment and the objective genes were analyzed for relative expression using the 2^-ΔΔCt^ method.

**Table 1.**
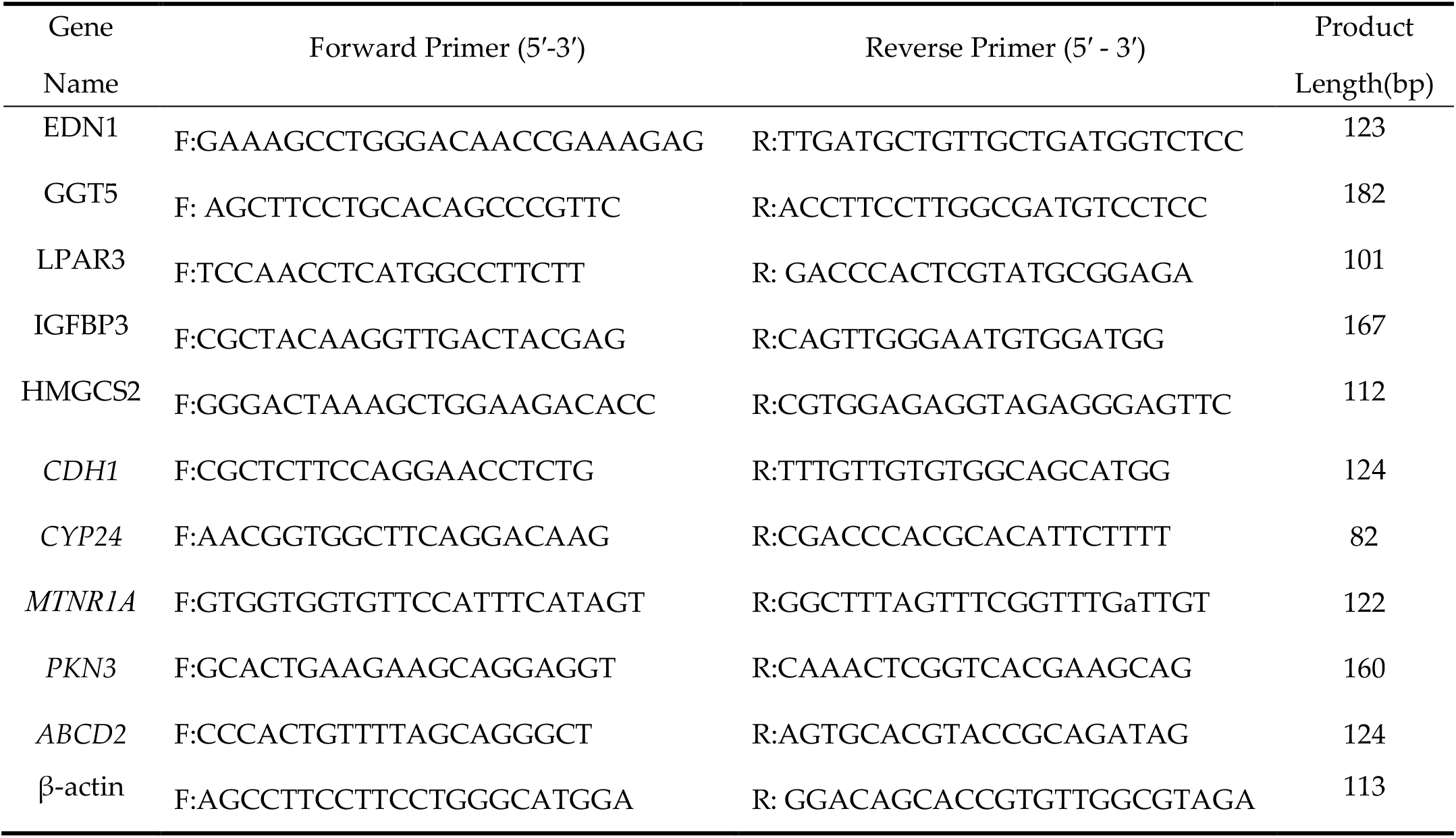
Primer information for *RT-*q*PCR*.

## 3. Results

### 3.1 Histological Observation of Testis

Figure 2 illustrates the histological analysis of testes of Tibetan sheep at high and low elevation. In the seminiferous tubules, there are Various spermatogenic cells and Sertoli cells and there is sperm production in the lumen. Comparison with low altitude Tibetan sheep, high altitude Tibetan sheep have more blood vessels between the seminiferous tubules. The measurement results (Table 2) indicate that diameter, area and thickness of the epithelium of the seminiferous tubules of high-altitude Tibetan sheep were significantly lower than those of low-altitude Tibetan sheep (P< 0.05). Compared to high elevation Tibetan sheep, significant increase in the areas of spermatogonia and primary spermatocytes at low-elevation Tibetan sheep (*P*< 0.05), number of Sertoli cells were much higher in Tibetan sheep at low-elevation (*P*< 0.05).

**Table 2.** Comparison with parameters of testicular seminiferous tubules in Tibetan sheep.

| Parameters | ZYH | ZYM |
| --- | --- | --- |
| Diameter of seminiferous tubules($\mu\text{m}$) | 97.34±0.50<sup>a</sup> | 118.32±0.51<sup>b</sup> |
| Seminiferous epithelium thickness($\mu\text{m}$) | 32.25±0.37<sup>a</sup> | 40.18±0.50<sup>b</sup> |
| Spermatogonia diameter ($\mu\text{m}$) | 5.76±0.45<sup>a</sup> | 8.76±0.40<sup>b</sup> |
| Primary spermatocyte diameter ($\mu\text{m}$) | 6.28±0.24<sup>a</sup> | 7.35±0.23<sup>b</sup> |
| Sperm cell diameter ($\mu\text{m}$) | 3.61±0.17<sup>a</sup> | 4.21±0.25<sup>b</sup> |
| Sertoli cell diameter ($\mu\text{m}$) | 6.17±0.28<sup>a</sup> | 6.74±0.40<sup>b</sup> |

Note: The table values are presented as mean ± SD. Varying alphabets in the alike line mean notable differences ( $P < 0.05$ ).

**Figure 1.**
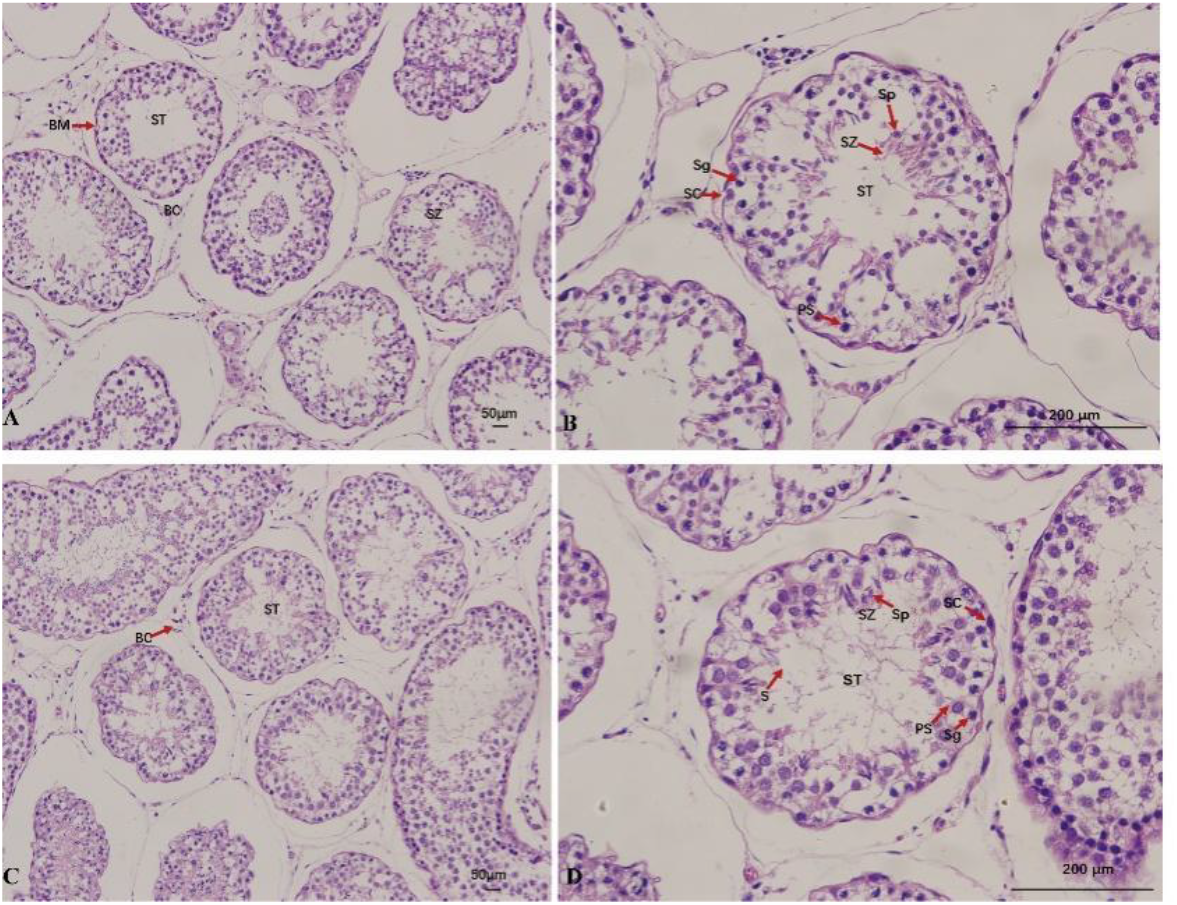
Histological analysis of testicular of Tibetan sheep in different elevations. (**A, B**): High-altitude Tibetan sheep testicular. (**C, D**): Low-altitude Tibetan sheep testicular. (**A, C**) stand for ×200 magnifying power. (**B, D**) stand for×400 magnifying power. ST: seminiferous tubules; BM: basement membrane; BC: blood capillary; Sc: Sertoli cells; Sg: spermatogonia; PS: primary spermatocytes; SZ: spermatozoa; Sp: Sperm cell;

**Figure 2.**
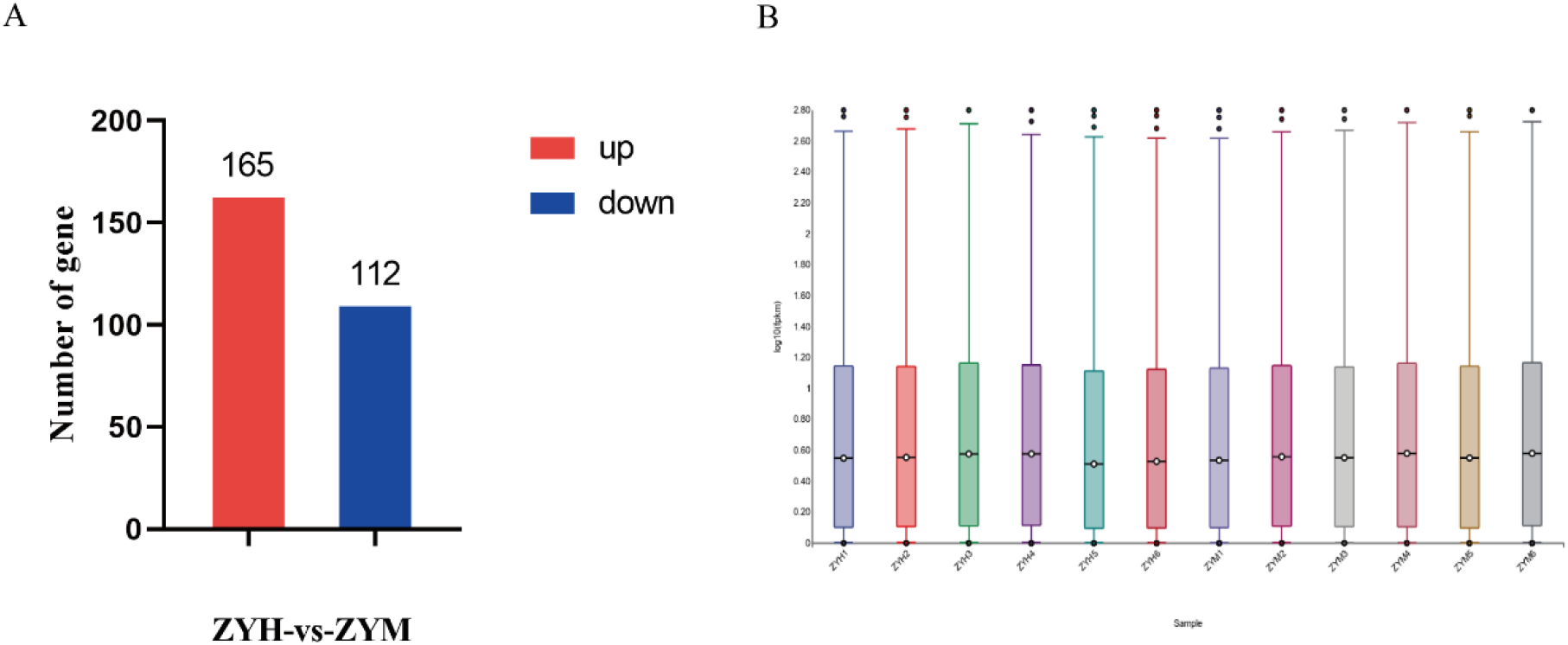
Distribution of gene expression levels. (A) Number of differentially expressed genes. (B) Volcano plot of the differentially expressed genes. ZYH: High altitude sheep; ZYM: Low altitude sheep

### 3.2 Summary of the RNA data of sheep testes

To screen for genes associated with testicular hypoxia adaptation, a total of twelve separate libraries were constructed and sequenced on the Illumina Hiseq platform. In Table 3, a summary of the information for these the RNA-Seq data are listed. In the lump, the twelve libraries generated 653,141,282 raw reads, after filtering the raw data, a total of 646,171,444 clean reads were remained. In the twelve samples, GC contents ranged from 53.87-56.07%, and which were in fulfilment of the basic rules of composition. Q20(%) ≥98.30%, Q30(%) ≥95.01%, which demonstrate the dependability of the RNA-seq figures and provide a reasonable basis for follow-up analyses. The clean data were matched to the reference genomes, the alignment results indicate that more than 94.30% of the reads were precisely calibrated with the sheep reference genomes and had higher matching rate. Among them, 3.15%-3.35% of clean read had multiple aligned position points, and 96.65%-96.82% of clean read had a single aligned position point, the latter can be used for further bioinformatic analysis. The overall gene expression level of each sample is further informed by comparing the FPKM values for each gene. This is shown in Figure 2B, the 12 samples showed similar in total gene expression levels.

**Table 3.** Summary of sequenced RNA-Seq data.

| Items | Raw reads | Clean reads | Q20(%) | Q30(%) | Total mapped | Multiple mapped | Unique mapped |
| --- | --- | --- | --- | --- | --- | --- | --- |
| ZYH1 | 580,430,32 | 574,735,68 | 98.67 | 96.03 | 562,284,29<br>(97.83%) | 184,147,8<br>(3.27%) | 543,869,51<br>(96.73%) |
| ZYH2 | 498,819,72 | 494,227,42 | 98.65 | 95.87 | 483,464,41<br>(97.82%) | 156,665,7<br>(3.24%) | 467,797,84<br>(96.76%) |
| ZYH3 | 504,663,38 | 499,358,68 | 98.47 | 95.41 | 486,484,56<br>(97.42%) | 153,8280<br>(3.16%) | 471,101,76<br>(96.84%) |
| ZYH4 | 480,660,00 | 475,737,56 | 98.55 | 95.63 | 465,213,83<br>(97.79%) | 146,543,1<br>(3.15%) | 450,559,52<br>(96.85%) |
| ZYH5 | 541,809,00 | 536,036,22 | 98.52 | 95.57 | 523,757,35<br>(97.71%) | 175,554,9<br>(3.35%) | 506,201,86<br>(96.65%) |
| ZYH6 | 570,682,06 | 565,283,12 | 98.7 | 96.08 | 552,702,36<br>(97.77%) | 185,067,6<br>(3.35%) | 534,195,60<br>(96.65%) |
| ZYM1 | 550,806,54 | 544,570,30 | 98.45 | 95.42 | 532,586,73<br>(97.80%) | 175,047,9<br>(3.29%) | 515,081,94<br>(96.71%) |
| ZYM2 | 577,825,24 | 571,769,42 | 98.6 | 95.83 | 559,202,41<br>(97.80%) | 181,873,5<br>(3.25%) | 541,015,06<br>(96.75%) |
| ZYM3 | 559,896,20 | 553,092,26 | 98.3 | 95 | 540,940,55<br>(97.80%) | 174,945,6<br>(3.23%) | 523,445,99<br>(96.77%) |
| ZYM4 | 526,462,10 | 520,999,68 | 98.58 | 95.74 | 509,146,26<br>(97.72%) | 164,768,6<br>(3.24%) | 492,669,40<br>(96.76%) |
| ZYM5 | 587,080,48 | 579,903,70 | 98.31 | 95.01 | 566,854,44<br>(97.75%) | 189,844,9<br>(3.35%) | 547,869,95<br>(96.65%) |
| ZYM6 | 552,277,78 | 546,000,40 | 98.34 | 95.03 | 514,852,36<br>(94.30%) | 169,651,5<br>(3.30%) | 497,887,21<br>(96.70%) |

### 3.3 Screening and Clustering Analysis of Differentially Expressed Genes

With FC ≥1.5, *P* <0.05 as the basis, to screen DEGs among libraries, we identified 277 DEGs between high and low-altitude Tibetan sheep, which 112 genes were down-regulated and 165 genes were up-regulated (Figure 2A). To gain a better understanding of differences in gene expression at high and low altitude Tibetan sheep, to cluster analysis of DEGs. The results revealed the DEGSs of the high-elevation Tibetan sheep flocks and the low elevation Tibetan sheep flocks were divided into two groups (Figure 3B). In addition, *IGFBP3* expression levels in the two groups showed an opposite tendency of the number of spermatogonia and Sertoli cells (Figure 4).

**Figure 3.**
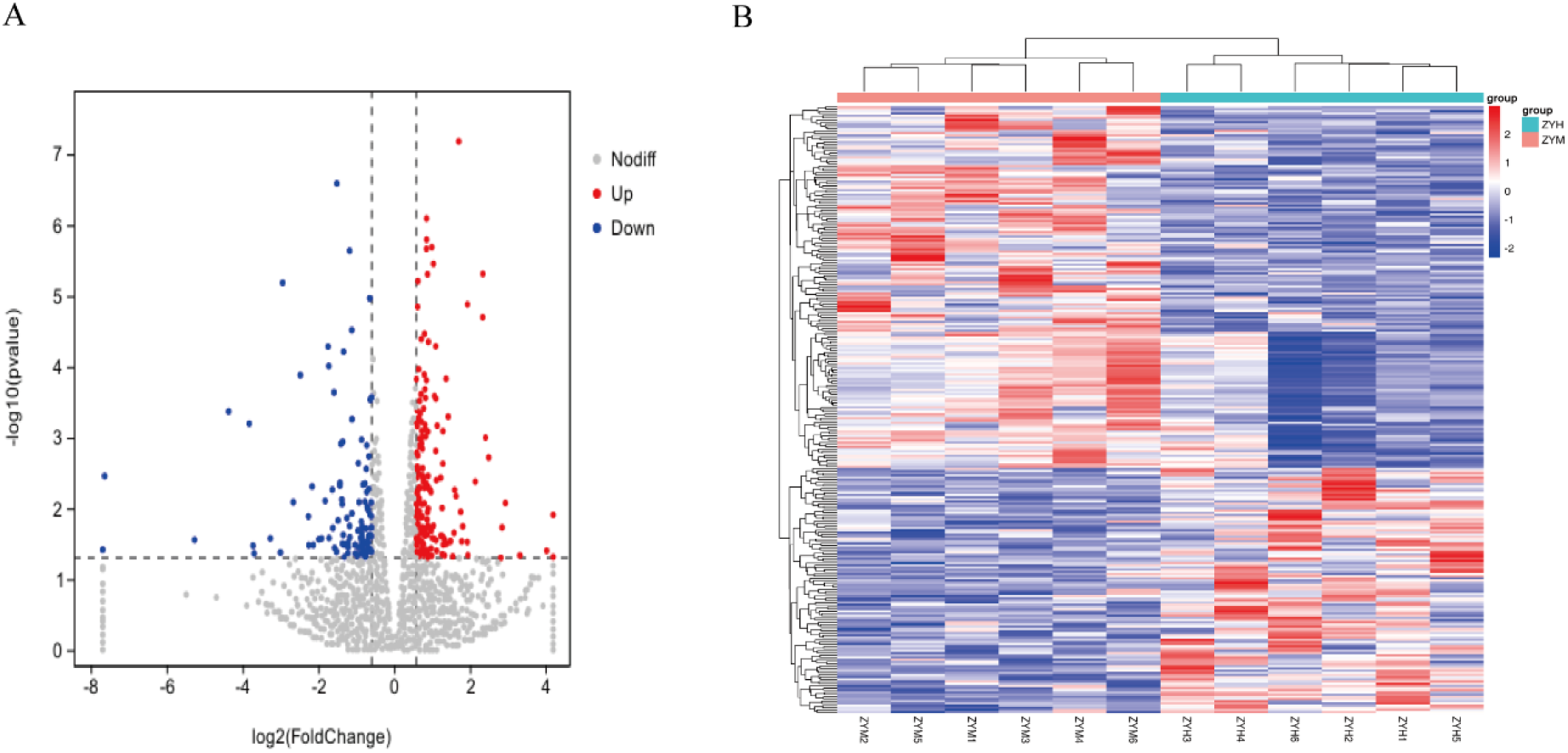
Differences in gene expression between high and low altitude Tibetan sheep populations. (**A**) Volcano diagram of DEGs. (**B**) Heat map of clustering analysis of DEGs. ZYH: High altitude Tibetan sheep; ZYM: Low-altitude Tibetan sheep.

**Figure 4.**
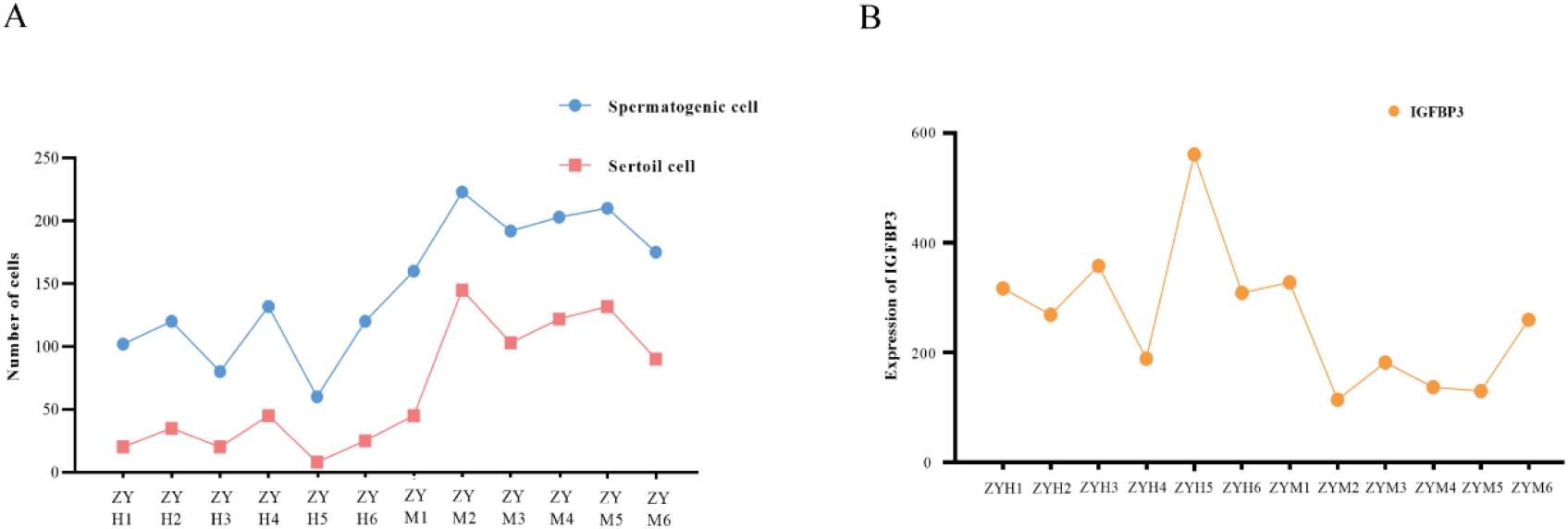
Expression levels of *IGFBP3* in samples and association with Sertoli cells and spermatogonia. (**A**) Number of Sertoli cells and spermatogonia. (**B**)Expression levels of *IGFBP3*.

### 3.4. GO and KEGG pathway Enrichment Analysis of DEGs

According to the GO enrichment analysis to grope for the functions of the DEGs in testicular adaptability,A total of 46 GO terms were remarkably enriched across the ZYH vs ZYM, including 20 biological processes, 11 molecular functions, and 15 cellular components. Biological processes which were mainly focused on “response to stimulus” (68 genes), “multicellular organismal process” (55 genes), and “regulation of multicellular organismal process” (31genes); cellular components which were mainly concentrated in “intrinsic component of membrane” (65 genes), and “integral component of membrane” (62genes); molecular functions which were mainly fastened on “cation binding” (35 genes), “metal ion binding” (32genes, Figure 5A). KEGG pathway enrichment analysis discovered that 70 pathways were enriched in ZYH vs ZYM, Figure 5B shows the top 20 pathways that are remarkably enriched in KEGG pathways. The results indicated that it was enriched in “Neuroactive ligand-receptor interaction” (14 genes), “Arachidonic acid metabolism” (5 genes), “Renin-angiotensin system” (3 genes). Besides, some pathways particularly associated with High-altitude environmental adaptation including HIF-1 signaling pathway (*EDN1*), P53 signaling pathway (*IGFBP3*), Glutathione metabolism (*GGT5*) and PPAR signaling pathway (*APOC3, PKC1*).

**Figure 5.**
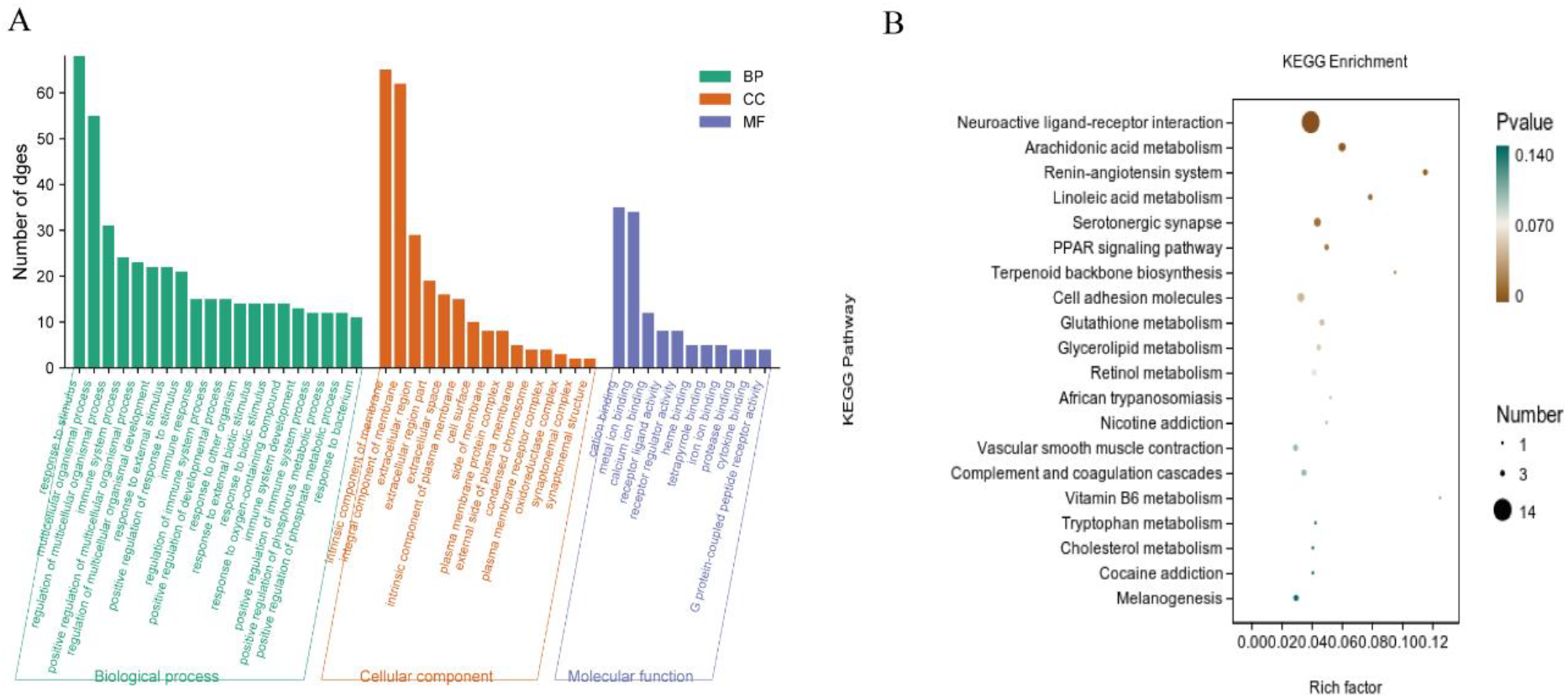
Functional analysis of DEGs between the ZYH and ZYM. (**A**) GO annotation of DEGs. (**B**) KEGG enrichment analysis pathway of DEGs

### 3.5. Validation of RNA-Seq data using RT-qPCR

In order to validate RNA-Seq data, ten genes were chosen for RT-qPCR analysis in Figure 6, The results revealed their expression patterns differ to some extent, but their expression patterns are exactly the same, thus indicating that the RNA-Seq data are credible.

**Figure 6.**
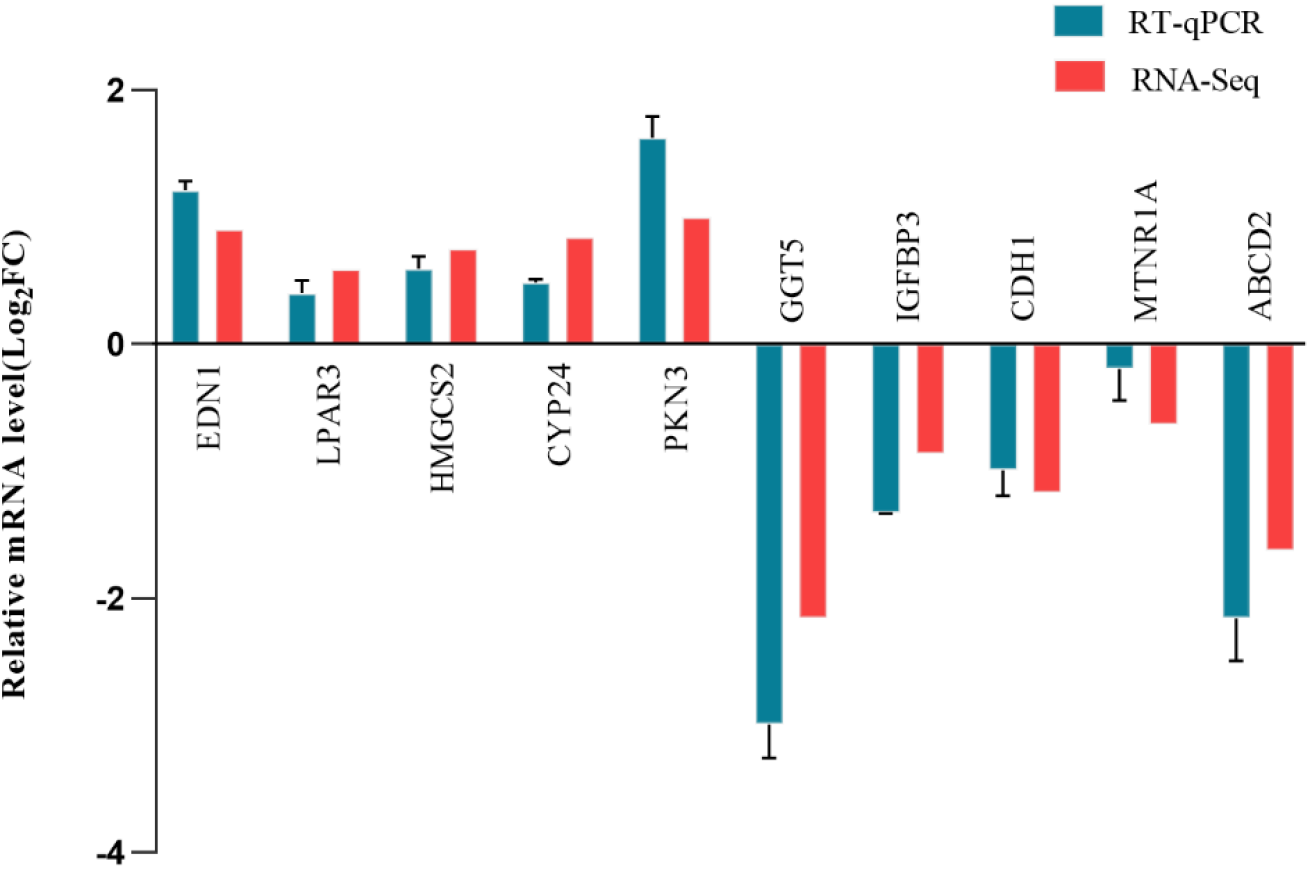
Validation of DEGs in testis of Tibetan sheep at high and low altitudes by RT-Qpcr

## 4. Discussion

Adaptation is taken as a core procedure of biological evolution, Phenotypic alterations in plateau adaptation are frequently accompanied by genetic modifications. Past investigations predominantly revolved around the former, on the comprehensive analysis of both fields being relatively scarce. RNA-Seq is a method that it is high-resolution, efficient, and deep-coverage to investigate the gene expression profiles and help to reveal the adaptive mechanism of the testis in the plateau environment. Analysis of histology architecture and gene expression of Tibetan sheep testis provides a full understanding for the mechanisms that regulate altitude differences. We select twelve Tibetan sheep for the study, RNA-Seq was used to screen key genes and pathways affecting altitude acclimation from them. On RNA-Seq analysis of twelve libraries of high and low altitude Tibetan sheep, 277 differentially expressed genes were screened out. Functional enrichment analysis yielded candidate signaling pathways which were explicitly enriched in Neuroactive ligand-receptor interaction (*LPAR3*), Arachidonic acid metabolism (*CYP2C19*), Renin-angiotensin system (*AGTR2*), P53 signaling pathway (*IGFBP3*), HIF-1 signaling pathway (*EDN1*), Glutathione metabolism (*GGT5*) and PPAR signaling pathway (*APOC3, PKC1*). In the KEGG analysis, most significantly enriched pathways is Neuroactive ligand-receptor interaction pathways, Relevant studies have shown that it could regulate reproductive activity and sexual maturity[18], genes related to Neuroactive ligand-receptor interaction are highly expressed in the testes of poultry[19]. In our analysis *LPAR3* genes were significantly up regulated in the low altitude Tibetan sheep group, *LPAR3* could interact with progesterone[20]and estrogen [21], thereby affecting reproductive performance. Study found that plasma estradiol levels were not altered in chronic hypoxia conditions, but progesterone levels were significantly reduced in In a hypoxic environment[22]. Therefore, we speculate that this pathway may adapt to its environment by regulating its reproductive activity, sexual maturation and hormonal interactions. Arachidonic acids, a complex secreted in an FSH-mediated way by Sertoli cells, which is one part of a cell-cell signaling modulatory network during spermatogenesis, and can induce spermatozoa and spermatocytes to release Ca2+ from internal stores[23]. Meanwhile, related studies have found that the overgrowth in Arachidonic acids metabolic pathway after hypoxia exposure may bring about maladjustment to hypoxia stimuli[24]. Cytochrome P450 2C19(*CYP2C19*) is a highly polymorphic gene[25], study found significant increase expression of the activity of *CYP2C19* in rats in the environment of acute high-altitude hypoxia[26]. In our study, *CYP2C19* genes Significantly enriched with Arachidonic acids metabolism pathway and significantly up regulated in the high-altitude Tibetan sheep group. Therefore, we hypothesized that this pathway may adapt to the oxygen environment through expression of *CYP2C19* activity and regulation of Ca2+ release in spermatocytes during spermatogenesis. The Renin-angiotensin system(RAS) pathway is involved in regulating vascular tone, also regulating vascular levels through vasoconstriction and regulating blood pressure by affecting plasma volume[27]. Besides, genes (*AGTR2*), up-regulated in high altitude Tibetan sheep, are associated not only with the regulation of blood pressure and blood circulation , but also with sperm capacitation and acrosome response[28]. Histological observation indicated that compared with low-altitude Tibetan sheep, high-altitude Tibetan sheep have more and blood vessels between the seminiferous tubules. Thus, our results infer that high altitude Tibetan sheep may enhanced testicular blood circulation using down-regulating these hypoxia response genes.

We found that some pathways were evidently enriched, for instance, Glutathione metabolism, and PPAR signaling pathway, P53 signaling pathway and HIF-1 signaling pathway. Glutathione is one of the most abundant tripeptides in tissues and cells , which plays an essential role in numerous cellular processes and adjusts significant cellular processes, including microtubule-related processes, cell growth, cell proliferation and immune responses[29]. As report goes, significant decreases in Glutathione(GSH) contents were observed in the muscles and blood of hypoxia-exposed rats during exposure to high altitude hypoxia[30]. *GGT5* is a key metabolic component that catalyzes the important antioxidant glutathione (GSH), which is expressed in poor levels in mammalian testicular interstitial cells (LCs). Over-expression of *GGT5* induced the expression of heme oxygenase 1 (HO-1), which inhibiting the activity of cytochrome P450 monooxygenase seriously impaired steroidogenesis of testicular [31]. In our study, *GGT5* genes were significantly downregulated in low altitude Tibetan sheep and enriched in the Glutathione metabolism pathway, Therefore, differential expression of *GGT5* may contribute to adaptability difference in sheep.

Peroxisome proliferator-activated receptors(PPAR) regulate fatty acid metabolism, glucose and energy metabolism[32], free fatty acids and glucose are necessary substrates to produce heat for cold-exposed animals[33–35], In our research, *PCK1* and *APOC3* enriched in the PPAR pathway. In addition , *PCK1*(phosphoenolpyruvate carboxykinase1) increases triglyceride production , and *APOC3* (apolipoprotein C3) is participated in the transport and uptake of free fatty acids[36]. High-density lipoprotein(HDL) is synthesized by a variety of multiple apoproteins such as *APOC3*, and are responsible for transporting cholesterol[37], HDL transmits cholesterol mostly to the steroidogenic organs(testes). To some extent, the production of testosterone depends on HDL-mediated cholesterol transport[38]. Therefore, the differential expression of *PKC1* and *APOC3* may be one of the reasons why Tibetan sheep are well adapted to the low-oxygen environment.

Hypoxia-inducible factor (HIF)-1, a major and key factor mediating the mammalian hypoxia response, regulates the oxygen balance at the cellular, tissue, and biological levels[39]. Recent studies have shown that HIF-1 signaling pathway, which responds to hypoxic conditions, plays an important role in angiogenesis[40]. Endothelin is a peptide family containing three isoforms, *EDN1, EDN2* and *EDN3*[41], whose *EDN1* can regulate vasoconstriction, affect cell proliferation, differentiation, migration, and apoptosis[42-43]. In this study, *EDN1* enriched in HIF-1 signaling pathway. Accordingly, we conjecture that Tibetan sheep adapts to the low-oxygen environment through angiogenesis and constriction in this pathway.

Insulin-like growth factor binding protein 3 (*IGFBP3*), a vital element in the *IGFBP* family, plays an imperative biological role in the regulation of apoptosis [44]. Long-term hypoxia exposure can promote *IGFBP3* protein synthesis and induce its secretion and promote apoptosis [45]. In p53 signaling pathway, *IGFBP3* genes enriched and evidently downregulated in the ZYM. Histological observations put up that the area of primary spermatocytes and spermatogonia in ZYM was obviously larger than ZYH. There were fewer Sertoli cells and stromal cells of ZYH. In all cases, expression levels of the *IGFBP3* gene showed an opposite tendency to the number of spermatogonia and Sertoli cells (Figure 6).In the seminiferous tubules, studies found the diameter, thickness and area of the epithelial cells of the seminiferous tubules were relevant for the number of spermatogenic cells[46]. We discovered the diameter, area and epithelial thickness were significantly higher in ZYM than in ZYH. Consequently, we hypothesized exorbitant apoptosis of germ cells of Tibetan sheep testis at high altitude may be one of the important causes of hypoxic acclimation.

## 5. Conclusions

To sum up, we analyzed testicular tissue of ZYH and ZYM. Spermatogonia cell counts were significantly higher in low altitude Tibetan sheep. Several genes (including *GGT5, AGTR2, EDN1, LPAR3, CYP2C19, IGFBP3, APOC3* and *PKC1*) were remarkably enriched in several pathways and terms, which may impact the Plateau adaptability of sheep by adjusting its reproductive activity and sexual maturation, and protecting Sertoli cells, various spermatocytes and spermatogenesis processes. The study offers valid transcriptomic references for altitude adaptation in Tibetan sheep at different elevation, provides more fundamental information on the regulatory mechanisms for improving testicular adaptation to the elevation environment in Tibetan sheep.

## Data availability

Reagents are available upon request. The authors affirm that all data necessary for confirming the conclusions of this article are represented fully within the article and its tables and figures.

## Author Contributions

B.Y. and X.W.: conception and design of the study. Y.Y. and Y.L.: analysis and interpretation of data. B.Y, R.G and M.M.: prepared the manuscript. All authors have read and agreed to the published version of the manuscript.

## Institutional Review Board Statement

The Ethics Committee of Gansu Agricultural University approved all the animal procedures (Approval number 2019-044).

## Data Availability Statement

All the raw data obtained in this study were deposited in the NCBI Sequence Read Archive (SRA) under the Bioproject accession number PRJNA10718.

## References

1. Wang, T.; Guo, Y.; Liu, S.; Zhang, C.; Cui, T.; Ding, K.; Wang, P.; Wang, X.; Z, Wang. KLF4, a Key Regulator of a Transitive Triplet, Acts on the TGF-β Signaling Pathway and Contributes to High-Altitude Adaptation of Tibetan Pigs. Front. Genet. 2021, 12, 628192. 10.3389/fgene.2021.628192.

2. Bai, L.; Liu, B.; Ji, C.; Zhao, S.; Liu, S.; Wang, R.; Wang, W.; Yao, P.; Li, X.; Fu, X.; Yu, H.; Liu, M.; Han, F.; Guan, N.; Liu, H.; Liu, D.; Tao, Y.; Wang, Z.; Yan, S.; Florant, G.; Butcher, M. T.; Zhang, J.; Zheng, H.; Fan, J.; E, Liu. Hypoxic and Cold Adaptation Insights from the Himalayan Marmot Genome. iScience 2019, 11, 505–507. 10.1016/j.isci.2019.01.019.

3. Ding, D.; Liu, G.; Hou, L.; Gui, W.; Chen, B.; L, Kang. Genetic Variation in PTPN1 Contributes to Metabolic Adaptation to High-Altitude Hypoxia in Tibetan Migratory Locusts. Nat. Commun. 2018, 9 (1), 4991. 10.1038/s41467-018-07529-8.

4. Fu, X.; Yang, Y.; Yan, Z.; Liu, M.; X, Wang. Transcriptomic Study of Spermatogenesis in the Testis of Hu Sheep and Tibetan Sheep. Genes 2022, 13 (12), 2212. 10.3390/genes13122212.

5. Wu, T.; B, Kayser. High Altitude Adaptation in Tibetans. High Alt. Med. Biol. 2006, 7 (3), 193– 208. 10.1089/ham.2006.7.193.

6. Sun, S.; Chen, Y.; R, Hu. Aquatic Hypoxia Disturbs Oriental River Prawn (Macrobrachium Nipponense) Testicular Development: A Cross-Generational Study. Environ. Pollut. 2020, 266, 115093. 10.1016/j.envpol.2020.115093.

7. Okumura, A.; Fuse, H.; Kawauchi, Y.; Mizuno, I.; T, Akashi. Changes in Male Reproductive Function after High Altitude Mountaineering. High Alt. Med. Biol. 2003, 4 (3), 349–353. 10.1089/152702903769192304.

8. Saxena, D. K. Effect of Hypoxia by Intermittent Altitude Exposure on Semen Characteristics and Testicular Morphology of Male Rhesus Monkeys. Int. J. Biometeorol. 1995, 38 (3), 137–140. 10.1007/BF01208490.

9. He, T.; Guo, H.; Shen, X.; Wu, X.; Xia, L.; Jiang, X.; Xu, Y.; Chen, D.; Zhang, Y.; Tan, D.; Y, Tan. Hypoxia-Induced Alteration of RNA Modifications in the Mouse Testis and Sperm. Biol. Reprod. 2021, 105 (5), 1171–1178. 10.1093/biolre/ioab142.

10. Farías, J. G.; Bustos-Obregón, E. J;, Reyes. G. Increase in Testicular Temperature and Vascularization Induced by Hypobaric Hypoxia in Rats. J. Androl. 2005, 26 (6), 693–697. 10.2164/jandrol.05013.

11. Hu, X.-J.; Yang, J.; Xie, X.-L.; Lv, F.-H.; Cao, Y.-H.; Li, W.-R.; Liu, M.-J.; Wang, Y.-T.; Li, J.-Q.; Liu, Y.-G.; Ren, Y.-L.; Shen, Z.-Q.; Wang, F.; Hehua, Ee.; Han, J.-L.; Li, M.-H. The Genome Landscape of Tibetan Sheep Reveals Adaptive Introgression from Argali and the History of Early Human Settlements on the Qinghai–Tibetan Plateau. Mol. Biol. Evol. 2019, 36 (2), 283–303. 10.1093/molbev/msy208.

12. Li, W.; Zeng, W.; Jin, X.; Xu, H.; Fang, X.; Ma, Z.; Cao, G.; Li, R.; L, Ma. High-Altitude Stress Orchestrates mRNA Expression and Alternative Splicing of Ovarian Follicle Development Genes in Tibetan Sheep. Animals 2022, 12 (20), 2812. 10.3390/ani12202812.

13. Zhao, P.; Li, S.; He, Z.; Zhao, F.; Wang, J.; Liu, X.; Li, M.; Hu, J.; Zhao, Z.; Y, Luo. Physiology and Proteomic Basis of Lung Adaptation to High-Altitude Hypoxia in Tibetan Sheep. Animals 2022, 12 (16), 2134. 10.3390/ani12162134.

14. Wang, F.; Liu, J.; Zeng, Q.; D, Zhuoga. Comparative Analysis of Long Noncoding RNA and mRNA Expression Provides Insights into Adaptation to Hypoxia in Tibetan Sheep. Sci. Rep. 2022, 12 (1), 6597. 10.1038/s41598-022-08625-y.

15. Kim, D.; Langmead, B. S;, Salzberg. L. HISAT: A Fast Spliced Aligner with Low Memory Requirements. Nat. Methods 2015, 12 (4), 357–360. 10.1038/nmeth.3317.

16. Pertea, M.; Pertea, G. M.; Antonescu, C. M.; Chang, T.-C.; Mendell, J. T. S;, Salzberg. L. StringTie Enables Improved Reconstruction of a Transcriptome from RNA-Seq Reads. Nat. Biotechnol. 2015, 33 (3), 290–295. 10.1038/nbt.3122.

17. Florea, L.; Song, L. S;, Salzberg. L. Thousands of Exon Skipping Events Differentiate among Splicing Patterns in Sixteen Human Tissues. F1000Research 2013, 2, 188. 10.12688/f1000research.2-188.v2.

18. Chen, B.; Liang, G.; Zhu, X.; Tan, Y.; Xu, J.; Wu, H.; Mao, H.; Zhang, Y.; Chen, J.; Rao, Y.; Zhou, M.; S, Liu. Gene Expression Profiling in Ovaries and Association Analyses Reveal HEP21 as a Candidate Gene for Sexual Maturity in Chickens. Animals 2020, 10 (2), 181. 10.3390/ani10020181.

19. Tang, B.; Hu, S.; Ouyang, Q.; Wu, T.; Lu, Y.; Hu, J.; Hu, B.; Li, L.; J, Wang. Comparative Transcriptome Analysis Identifies Crucial Candidate Genes and Pathways in the Hypothalamic-Pituitary-Gonadal Axis during External Genitalia Development of Male Geese. BMC Genomics 2022, 23 (1), 136. 10.1186/s12864-022-08374-2.

20. Liszewska, E.; Reinaud, P.; Dubois, O.; G, Charpigny. Lysophosphatidic Acid Receptors in Ovine Uterus during Estrous Cycle and Early Pregnancy and Their Regulation by Progesterone. Domest. Anim. Endocrinol. 2012, 42 (1), 31–42. 10.1016/j.domaniend.2011.08.003.

21. Diao, H.; Li, R.; El Zowalaty, A. E.; Xiao, S.; Zhao, F.; Dudley, E. A.; X, Ye. Deletion of Lysophosphatidic Acid Receptor 3 (Lpar3) Disrupts Fine Local Balance of Progesterone and Estrogen Signaling in Mouse Uterus During Implantation1. Biol. Reprod. 2015, 93 (5). 10.1095/biolreprod.115.131110.

22. Shaw, S.; Kumar, U.; Bhaumik, G.; Reddy, M. P. K.; Kumar, B.; D, Ghosh. Alterations of Estrous Cycle, 3β Hydroxysteroid Dehydrogenase Activity and Progesterone Synthesis in Female Rats after Exposure to Hypobaric Hypoxia. Sci. Rep. 2020, 10 (1), 3458. 10.1038/s41598-020-60201-4.

23. Paillamanque, J.; Sanchez-Tusie, A.; Carmona, E. M.; Treviño, C. L.; Sandoval, C.; Nualart, F.; Osses, N. J;, Reyes. G. Arachidonic Acid Triggers [Ca2+]i Increases in Rat Round Spermatids by a Likely GPR Activation, ERK Signalling and ER/Acidic Compartments Ca2+ Release. PLOS ONE 2017, 12 (2), e0172128. 10.1371/journal.pone.0172128.

24. Liu, C.; Liu, B.; Liu, L.; Zhang, E.-L.; Sun, B.; Xu, G.; Chen, J.; Y, Gao. Arachidonic Acid Metabolism Pathway Is Not Only Dominant in Metabolic Modulation but Associated With Phenotypic Variation After Acute Hypoxia Exposure. Front. Physiol. 2018, 9, 236. 10.3389/fphys.2018.00236.

25. Jin, T.; Zhang, X.; Geng, T.; Shi, X.; Wang, L.; Yuan, D.; L, Kang. Genotype-Phenotype Analysis of CYP2C19 in the Tibetan Population and Its Potential Clinical Implications in Drug Therapy. Mol. Med. Rep. 2016, 13 (3), 2117–2123. 10.3892/mmr.2016.4776.

26. Li, X.; Wang, X.; Li, Y.; Yuan, M.; Zhu, J.; Su, X.; Yao, X.; Fan, X.; Y, Duan. Effect of Exposure to Acute and Chronic High-Altitude Hypoxia on the Activity and Expression of CYP1A2, CYP2D6, CYP2C9, CYP2C19 and NAT2 in Rats. Pharmacology 2014, 93 (1–2), 76–83. 10.1159/000358128.

27. Rupert, J. L.; Kidd, K. K.; Norman, L. E.; Monsalve, M. V.; Hochachka, P. W. D;, Devine. V. Genetic Polymorphisms in the Renin-Angiotensin System in High-Altitude and Low-Altitude Native American Populations. Ann. Hum. Genet. 2003, 67 (1), 17–25. 10.1046/j.1469-1809.2003.00004.x.

28. Corda, P. O.; Santiago, J.; M, Fardilha. G-Protein Coupled Receptors in Human Sperm: An In Silico Approach to Identify Potential Modulatory Targets. Molecules 2022, 27 (19), 6503. 10.3390/molecules27196503.

29. Wang, L.; Ahn, Y. J.; R, Asmis. Sexual Dimorphism in Glutathione Metabolism and Glutathione-Dependent Responses. Redox Biol. 2020, 31, 101410. 10.1016/j.redox.2019.101410.

30. Singh, S. N.; Vats, P.; Kumria, M. M. L.; Ranganathan, S.; Shyam, R.; Arora, M. P.; Jain, C. L.; K, Sridharan. Effect of High Altitude (7,620 m) Exposure on Glutathione and Related Metabolism in Rats. Eur. J. Appl. Physiol. 2001, 84 (3), 233–237. 10.1007/s004210170010.

31. Li, W.; Wu, Z.; Zhang, S.; Cao, R.; Zhao, J.; Sun, Z.; W, Zou. Augmented Expression of Gamma-Glutamyl Transferase 5 (GGT5) Impairs Testicular Steroidogenesis by Deregulating Local Oxidative Stress. Cell Tissue Res. 2016, 366 (2), 467–481. 10.1007/s00441-016-2458-y.

32. Nakamura, M. T.; Yudell, B. E. J;, Loor. J. Regulation of Energy Metabolism by Long-Chain Fatty Acids. Prog. Lipid Res. 2014, 53, 124–144. 10.1016/j.plipres.2013.12.001.

33. Paul, P. W;, Holmes. L. Free Fatty Acid Metabolism during Stress: Exercise, Acute Cold Exposure, and Anaphylactic Shock. Lipids 1973, 8 (3), 142–150. 10.1007/BF02531811.

34. Smith, O. L. S;, Davidson. B. Shivering Thermogenesis and Glucose Uptake by Muscles of Normal or Diabetic Rats. Am. J. Physiol.-Regul. Integr. Comp. Physiol. 1982, 242 (1), R109–R115.10.1152/ajpregu.1982.242.1.R109.

35. Sepa-Kishi, D. M.; Sotoudeh-Nia, Y.; Iqbal, A.; Bikopoulos, G. R;, Ceddia. B. Cold Acclimation Causes Fiber Type-Specific Responses in Glucose and Fat Metabolism in Rat Skeletal Muscles. Sci. Rep. 2017, 7 (1), 15430. 10.1038/s41598-017-15842-3.

36. Zhang, Z.; Sui, Z.; Zhang, J.; Li, Q.; Zhang, Y.; F, Xing. Transcriptome Sequencing-Based Mining of Genes Associated With Pubertal Initiation in Dolang Sheep. Front. Genet. 2022, 13, 818810. 10.3389/fgene.2022.818810.

37. Wayne Hou, J.; Collins, D. C. R;, Schleicher. L. Sources of Cholesterol for Testosterone Biosynthesis in Murine Leydig Cells*. Endocrinology 1990, 127 (5), 2047–2055. 10.1210/endo-127-5-2047.

38. Yang, L.; Ma, T.; Zhao, L.; Jiang, H.; Zhang, J.; Liu, D.; Zhang, L.; Wang, X.; Pan, T.; Zhang, H.; Wang, A.; Chao, H.-W.; Jin, Y.; H, Chen. Circadian Regulation of Apolipoprotein Gene Expression Affects Testosterone Production in Mouse Testis. Theriogenology 2021, 174, 9–19. 10.1016/j.theriogenology.2021.06.023.

39. Devraj, G.; Beerlage, C.; Brüne, B. V;, Kempf. A.J. Hypoxia and HIF-1 Activation in Bacterial Infections. Microbes Infect. 2017, 19 (3), 144–156. 10.1016/j.micinf.2016.11.003.

40. Bai, H.; Guo, X.; Tan, Y.; Wang, Y.; Feng, J.; Lei, K.; Liu, X.; Xiao, Y.; C, Bao. Hypoxia Inducible Factor-1 Signaling Pathway in Macrophage Involved Angiogenesis in Materials-Instructed Osteo-Induction. J. Mater. Chem. B 2022, 10 (34), 6483–6495. 10.1039/D2TB00811D.

41. Thaete, L. G.; Jilling, T.; Synowiec, S.; Khan, S. M;, Neerhof. G. Expression of Endothelin 1 and Its Receptors in the Hypoxic Pregnant Rat1. Biol. Reprod. 2007, 77 (3), 526–532. 10.1095/biolreprod.107.061820.

42. Cervar, M.; Puerstner, P.; Kainer, F.; G, Desoye. Endothelin-1 Stimulates the Proliferation and Invasion of First Trimester Trophoblastic Cells in Vitro--a Possible Role in the Etiology of Pre-Eclampsia? J. Investig. Med. Off. Publ. Am. Fed. Clin. Res. 1996, 44 (8), 447–453.

43. Shichiri, M.; Kato, H.; Marumo, F.; Y, Hirata. Endothelin-1 as an Autocrine/Paracrine Apoptosis Survival Factor for Endothelial Cells. Hypertension 1997, 30 (5), 1198–1203. 10.1161/01.HYP.30.5.1198.

44. Butt, A. J. A;, Williams. C. [No Title Found]. APOPTOSIS 2001, 6 (3), 199–205. 10.1023/A:1011388710719.

45. Chang, R.-L.; Lin, J.-W.; Hsieh, D. J.-Y.; Yeh, Y.-L.; Shen, C.-Y.; Day, C.-H.; Ho, T.-J.; Viswanadha, V. P.; Kuo, W.-W.; Huang, C.-Y. Long-Term Hypoxia Exposure Enhanced IGFBP-3 Protein Synthesis and Secretion Resulting in Cell Apoptosis in H9c2 Myocardial Cells. Growth Factors 2015, 33 (4), 275–281. 10.3109/08977194.2015.1077824.

46. Youn, J. S.; Cha, S. H.; Park, C. W.; Yang, K. M.; Kim, J. Y.; Koong, M. K.; Kang, I. S.; Song, I. O. S;, Han. C. Predictive Value of Sperm Motility Characteristics Assessed by Computer-Assisted Sperm Analysis in Intrauterine Insemination with Superovulation in Couples with Unexplained Infertility. Clin. Exp. Reprod. Med. 2011, 38 (1), 47. 10.5653/cerm.2011.38.1.47.

